# The effect of stimulus envelope shape on the auditory steady-state response

**DOI:** 10.1101/541359

**Authors:** Jana Van Canneyt, Michael Hofmann, Jan Wouters, Tom Francart

**Author notes:** Email addresses (Jana Van Canneyt), (Michael Hofmann), (Jan Wouters), (Tom Francart).

## Abstract

Auditory steady-state responses (ASSRs) are auditory evoked potentials that reflect phase-locked neural activity to periodic stimuli. ASSRs are often evoked by tones with a modulated envelope, with sinusoidal envelopes being most common. However, it is unclear if and how the shape of the envelope affects ASSR responses. In this study, we used various trapezoidal modulated tones to evoke ASSRs (modulation frequency = 40 Hz) and studied the effect of four envelope parameters: attack time, hold time, decay time and off time. ASSR measurements in 20 normal hearing subjects showed that envelope shape significantly influenced responses: increased off time and/or increased decay time led to responses with a larger signal-to-noise-ratio (SNR). Response phase delay was significantly influenced by attack time and to a lesser degree by off time. We also simulated neural population responses that approximate ASSRs with a model of the auditory periphery (Bruce et al. 2018). The modulation depth of the simulated responses, i.e. the difference between maximum and minimum firing rate, correlated highly with the response SNRs found in the ASSR measurements. Longer decay time and off time enhanced the modulation depth both by decreasing the minimum firing rate and by increasing the maximum firing rate. In conclusion, custom envelopes with long decay and off time provide larger response SNRs and the benefit over the commonly used sinusoidal envelope was in the range of several dB.

## 1. Introduction

Auditory neurons are more likely to fire during the rising phases of the waveform of a perceived sound than during the falling phases. This phase-locking ability allows neurons to fire synchronously with each other and with (a delayed version of) the auditory input. Moreover, phase-locking occurs throughout the auditory pathway, from the primary auditory nerve fibres up to the auditory cortex. Using electroencephalogram (EEG) measurements, the compound synchronized neural firing activity of a large population of neurons can be registered from the scalp. As a result of phase-locking, the auditory neural response in the EEG signal shows the same periodicity as the evoking stimulus. This phenomenon underlies a group of auditory evoked potentials called frequency following responses. Stimuli can be designed such that the neurons predominantly phase-lock to the stimulus envelope and not to the temporal fine structure. This results in envelope following responses, a subcategory of frequency following responses. Auditory steady-state responses (ASSRs) are envelope following responses evoked by steadily repeated or modulated stimuli (Galambos et al., 1981; Picton et al., 2003). The discrete frequency components of the ASSR are constant in amplitude (Van Eeckhoutte et al., 2018) and phase over time, allowing the responses to be evaluated in the frequency domain.

ASSRs are valuable for auditory research because the responses reflect important characteristics of auditory processing, including hearing sensitivity (Rance et al., 1995), loudness growth (Van Eeckhoutte et al., 2016) and temporal modulation sensitivity (Luke et al., 2015). ASSR reponse strength is also correlated with more holistic perceptual performance, e.g. word recognition (Dimitrijevic et al., 2004), phoneme perception and speech in noise perception (Alaerts et al., 2009). Clinically, ASSRs are used to determine frequency specific hearing thresholds both in adults (Rance et al., 1995) and in children (Rance and Rickards, 2002; Luts et al., 2006; Van Maanen and Stapells, 2010). Even though not yet implemented, recent studies point towards other clinical applications of the ASSRs, e.g. the diagnosis of dyslexia (Poelmans et al., 2012; Vanvooren, 2015; De Vos et al., 2017) and the assessment of age-related changes in hearing that are not reflected in the audiogram (Goossens et al., 2016; Dimitrijevic et al., 2016).

For clinical practice, it is important that responses can be measured quickly in a noisy environment. Therefore, stimuli and measurement procedures are optimized to obtain responses with a large signal-to-noise ratio (SNR). The most commonly used stimuli to evoke ASSRs are sinusoidally amplitude modulated (SAM) stimuli, which consist of a carrier (pure tone, triangle wave, noise band, etc.) and a sinusoidal modulator. The effects of modulation frequency, carrier frequency and carrier type on the evoked ASSRs are well understood (Picton et al., 2003). By definition, SAM stimuli have a sinusoidal envelope. However, there is no evidence that this envelope shape is most optimal to evoke ASSRs. In fact, there is very little information available on the effect of envelope shape on the ASSR. The most relevant study by John et al. (2002) compared the strength of ASSRs evoked by exponential sine envelopes, i.e. *sin*^*N*^ (*t*), with different exponents. They found that responses were 20-30 % larger for exponential stimuli with N>1, than for SAM stimuli (N=1).

The effect of envelope shape is better understood when it concerns the perception of interaural time differences (ITDs), which are important for sound localisation. Just like ASSRs, ITD perception heavily depends on synchronous neural firing. Bernstein and Trahiotis (2002) compared sensitivity to ITDs with SAM stimuli and transposed tones. Transposed tones are pure tones modulated with a linearly half-wave rectified sinusoid and were first introduced by van de Par and Kohlrausch (1997). A higher sensitivity to ITDs was observed with transposed tones. In another study, Bernstein and Trahiotis (2009) investigated the effect of exponential sinusoidal modulation with different exponents on ITD perception. Similar to what John et al. (2002) found for ASSR, higher sensitivity to ITDs was obtained with larger exponents.

The aforementioned studies found better results with exponentially modulated or transposed stimuli than with SAM stimuli, suggesting that non-sinusoidal envelopes enhanced neural synchronisation. Physiological animal studies have confirmed this at neural level for transposed tones (Griffin, 2005; Dreyer and Delgutte, 2006). The enhancement is attributed to longer near-zero sections (”off time”) and steeper attacks in the envelope of these stimuli compared to SAM stimuli. A longer off time is believed to allow the neurons to recover more from refractory effects and to therefore increase firing capacity at the next attack. A steeper attack slope decreases the variability in first spike timing and also increases spike count in response to the attack (Heil, 1997a,b; Heil and Irvine, 1997).

To investigate more conclusively how envelope parameters like off time and attack slope affect neural synchrony, Klein-Hennig et al. (2011) employed a more controlled study design. They measured ITD perception for stimuli with a trapezoidal envelope consisting of four segments: attack, hold, decay and off. The effect of envelope shape was studied by varying the duration of each of the segments independently. Confirming results from the previous studies, they observed that the duration of the off and the attack influenced ITD sensitivity, while changing the duration of the hold and the decay had no significant effect. Similar results were found by Laback et al. (2011) and Greenberg et al. (2017).

The objective of this study was to investigate the effect of envelope shape on the ASSR. Using a study design similar to Klein-Hennig et al. (2011) and Laback et al. (2011), we manipulated envelope shape and observed the effect on the signal-to-noise-ratio (SNR) and the phase delay of the ASSR. Based on the available literature, we expected that shorter attack and longer off time would be associated with stronger responses. To obtain a deeper understanding, we studied the effect of envelope shape on neural firing in the auditory periphery using the model of Bruce et al. (2018).

## 2. ASSR measurements - material and methods

### 2.1. Subjects

Twenty young subjects (15 females, 5 males) participated in the ASSR measurements. Their ages ranged from 18 to 32 years old (mean = 23.9). All participants had normal hearing, which was confirmed using pure-tone audiometry (octave frequencies between 125 and 8000 Hz, all thresholds < 25 dB SPL). The experiments were approved by the medical ethics committee of the University Hospital of Leuven and all subjects signed an informed consent form before participating.

### 2.2. Conditions and stimuli

Stimuli were trapezoidally modulated 900 Hz pure tones (see figure 1). The trapezoidal envelope of each modulation period was based on the study of Klein-Hennig et al. (2011) and consisted of four segments: attack, hold, decay and off. The attack and decay were formed by cosine ramps between the desired start and end times. The hold and off had a constant amplitude of 1 and 0 respectively, resulting in a modulation depth of 100%. Envelope shape was manipulated by varying the duration of the four segments. Consequently, the four envelope parameters were attack time, hold time, decay time and off time.

**Figure 1:**
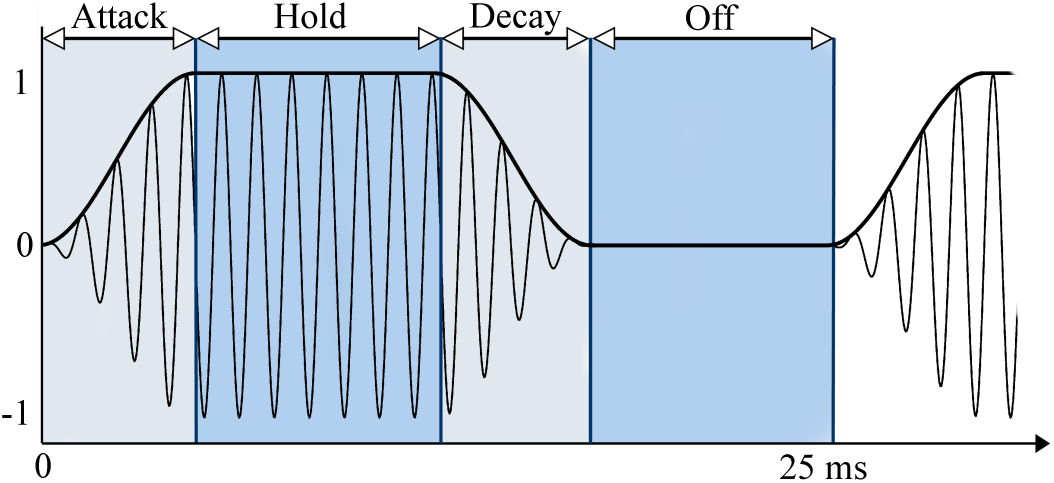
Example of a trapezoidally modulated stimulus. The carrier was a 900 Hz pure tone. The modulation frequency was 40 Hz, corresponding to a envelope period of 25 ms. The modulation envelope consisted of four segments: attack, hold, decay and off, which are indicated on the figure. The shape of the envelope was modified by varying the duration of the segments.

Varying one of the envelope parameters while keeping the others constant, alters the duration of the envelope period, which is equal to the sum of the four parameters. This results in a difference in modulation frequency over stimuli. Klein-Hennig et al. (2011) argued that ITD perception is not influenced by changes in modulation frequency and therefore accepted this variability in their study design. However, ASSRs are known to vary greatly in amplitude with modulation frequency (Gransier, 2018). For lower modulation frequencies (≤ 50 Hz), cortical sources contribute strongly to the ASSR. For higher modulation frequencies subcortical sources become more dominant (Herdman et al., 2002; Bidelman, 2018). This introduces a low-pass characteristic to the response amplitude, because subcortical responses are smaller than cortical responses when measured at the scalp. In addition, phase-locked responses from the different sources have different latencies. Through constructive and destructive phase interferences, the response at the scalp is attenuated or enhanced in a frequency-specific manner (Tichko and Skoe, 2017). To avoid confounding effects in this study, the modulation frequency had to remain constant. More specifically, all stimulus envelopes in the current study had an envelope period of 25 ms, corresponding to a modulation frequency of 40 Hz. This modulation frequency was chosen because it yields large ASSR responses (Picton et al., 2003).

To study the effect of the envelope parameters without changing the modulation frequency, two parameters were varied together, e.g. increasing attack time while decreasing hold time. The other two parameters were kept constant. All possible combinations of the four envelope parameters were included. This formed six conditions, which were named after the parameters that were varied: attack-hold, attack-decay, attack-off, hold-decay, hold-off and decay-off. The varied parameters generally took values between 0 and 16.25 ms, but the shortest attack and decay times were limited to 1.25 ms to avoid excessive spectral broadening of the stimuli (Klein-Hennig et al., 2011). When parameters were not varied, they had the constant values of 5, 5, 5, and 2.5 ms for attack time, hold time, decay time and off time, respectively. Every condition included 4 or 6 envelope shapes, i.e. two extremes and intermediate stages in between (see figure 2), summing up to a total of 24 unique envelope shapes (32 when duplicates are included). Besides these, a typical sinusoidal modulation, i.e. SAM stimulus, was added to the protocol, resulting in a total of 25 unique stimuli.

**Figure 2:**
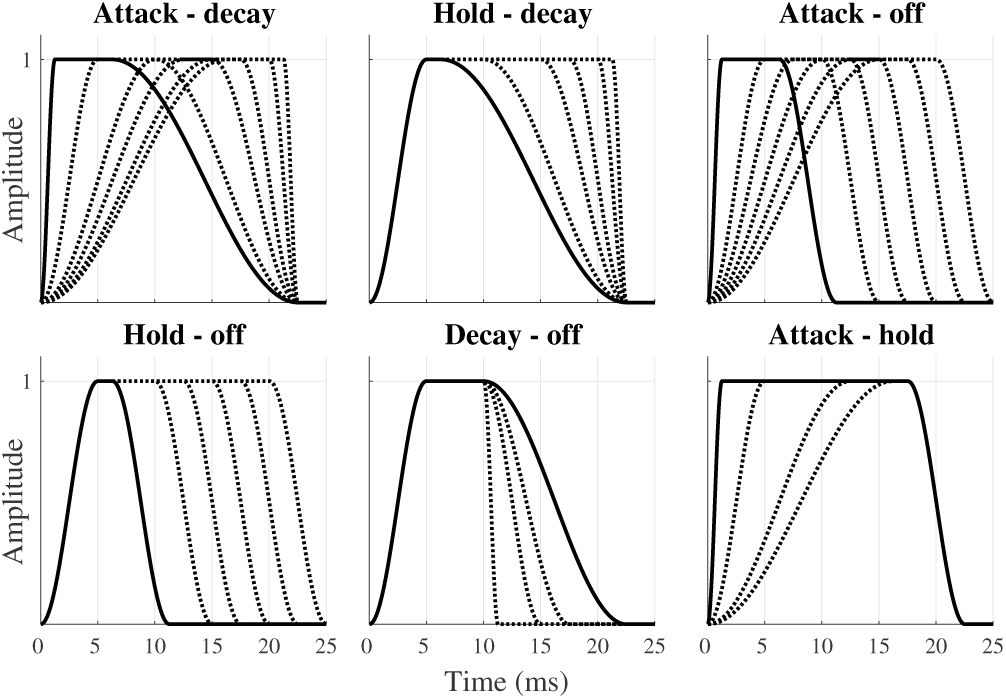
Visualisation of the conditions and the stimulus envelopes that represent them. Modulation frequency was equal to 40 Hz for all stimuli. Envelopes were defined between 0 and 1 resulting in 100 % modulation depth.

Stimuli were created in MATLAB R2016b (The MathWorks Inc., 2016) with amplitudes defined between −1 and 1 (32 bit) and a sampling frequency of 32 kHz. They were presented through an insert phone (3M, E-A-RTONE) in the right ear using custom developed software (Hofmann and Wouters, 2010). Each stimulus lasted 5 minutes and consisted of 300 repetitions of the same 1.024 second stimulus segment. The beginning of each repetition was marked with a trigger - which was used in the analysis - and a smooth transition between repetitions was ensured. All stimuli were presented at the same peak level, which was defined as the level required to present the SAM tone at 75 dB SPL. This calibration was done using a 2 cc artificial ear (Bruel & Kjær, type 4152) and a sound level meter (Bruel & Kjær, type 2250). The presentation order of the stimuli was randomized among subjects.

### 2.3. Response measurement

ASSR were measured with a 64-channel Biosemi ActiveTwo EEG recording system (sampling rate = 8192 Hz). The 64 Ag/AgCl active scalp electrodes were placed on a cap according to the international standardized 10-10 system (American Clinical Neurophysiology society, 2006). Two extra electrodes, CMS and DRL, functioned as the common electrode and the current return path, respectively. Subjects were seated in an electromagnetically-shielded soundproof booth and watched a silent movie with subtitles. This encouraged the subjects to maintain the same passive listening state throughout the 25 measurements which lasted about 2.5 hours in total.

### 2.4. Response analysis

The EEG signals were processed with MATLAB R2016b (The MathWorks Inc., 2016). We used 12 EEG electrodes located at the back of the head (CP5, CP6, P5, P6, PO3, PO4, O1, O2, PO7, PO8, TP7 and TP8) and referenced them to Cz, located at the top of the head. First, the signals of these 12 electrodes were averaged to reduce recording noise. Then, the averaged signal was high-pass filtered with a cut-off frequency of 2 Hz to remove any DC component (2nd order Butterworth filter). The filtered signal was divided into 300 epochs of 1.024 s based on the triggers from the stimulation software. The epochs with the 5% largest peak-to-peak amplitudes were removed to reduce the amount of (muscle) artefacts. Each of the remaining epochs was converted into a complex frequency spectrum using a Fast Fourier Transform with a frequency resolution of 0.97 Hz.

For every epoch, the amplitude and phase of the ASSR response was determined for the frequency bin corresponding to the modulation frequency, i.e. 40 Hz. The mean response amplitude and phase were determined by vector averaging over epochs. The noise level was defined as the variance over epochs. Response SNR was determined as the ratio between response amplitude and recording noise. Responses were statistically evaluated using the Hotelling *T* ^2^ test (Hotelling, 1931) which compares the response amplitude with the recording noise. A significance level of 5 % corresponds to an SNR of 4.8 dB SPL, calculated with the method of Dobie and Wilson (1996). Across all measurements, two did not yield significant ASSR responses and were excluded from further analysis. Apart from the SNR, we also investigated the effect of the envelope parameters on phase delay. Phase delay is the phase difference between the response and the evoking stimulus, that originates from the time delay introduced by neural processing. It is calculated by subtracting the average phase of the response from 360°.

The effects of the envelope parameters on response SNR and phase delay were visualised and analysed in R (R Core Team, 2018). Since ASSRs, and frequency following responses in general, are known to have large inter-subject variability (but small intra-subject variability), normalized response SNR and phase delays are shown in the figures. Normalization was done by subtracting the mean value for a subject from all the data points of that subject. In the figures, data is presented per condition, together with a linear regression line. For the statistical analysis of the effects across conditions, the non-normalized data was used. We employed a generalized linear mixed model with a random intercept per subject (package lme4, version 1.1.17, Douglas et al. (2015)). The random intercept accounts for the inter-subject variability. Significance of the model predictors was evaluated with a t-test (significance level = 0.05) using Satterthwaite’s method to estimate the degrees of freedom (Satterthwaite, 1946). Visual inspection of the residual plots from any of the models reported in this paper revealed no deviations from homoscedascity or normality.

## 3. ASSR measurements - results

### 3.1. Response SNR

The average amplitude of the ASSRs measured in this study was 272 nV (range = 34-784 nV) and the average amplitude of the recording noise was 24 nV (range = 14-37 nV). Response SNRs had an average value of 20.1 dB (range = 5.02 - 31.1 dB) which is well above the threshold for significance of 4.8 dB. The mean SNR per subject ranged between 14.2 dB and 27.1 dB, illustrating the large inter-individual differences and the need for normalization. The normalized SNRs for the six conditions are presented in figure 3 together with linear regression fits per condition. The median normalized response SNR for the SAM stimulus is indicated with the dashed line.

**Figure 3:**
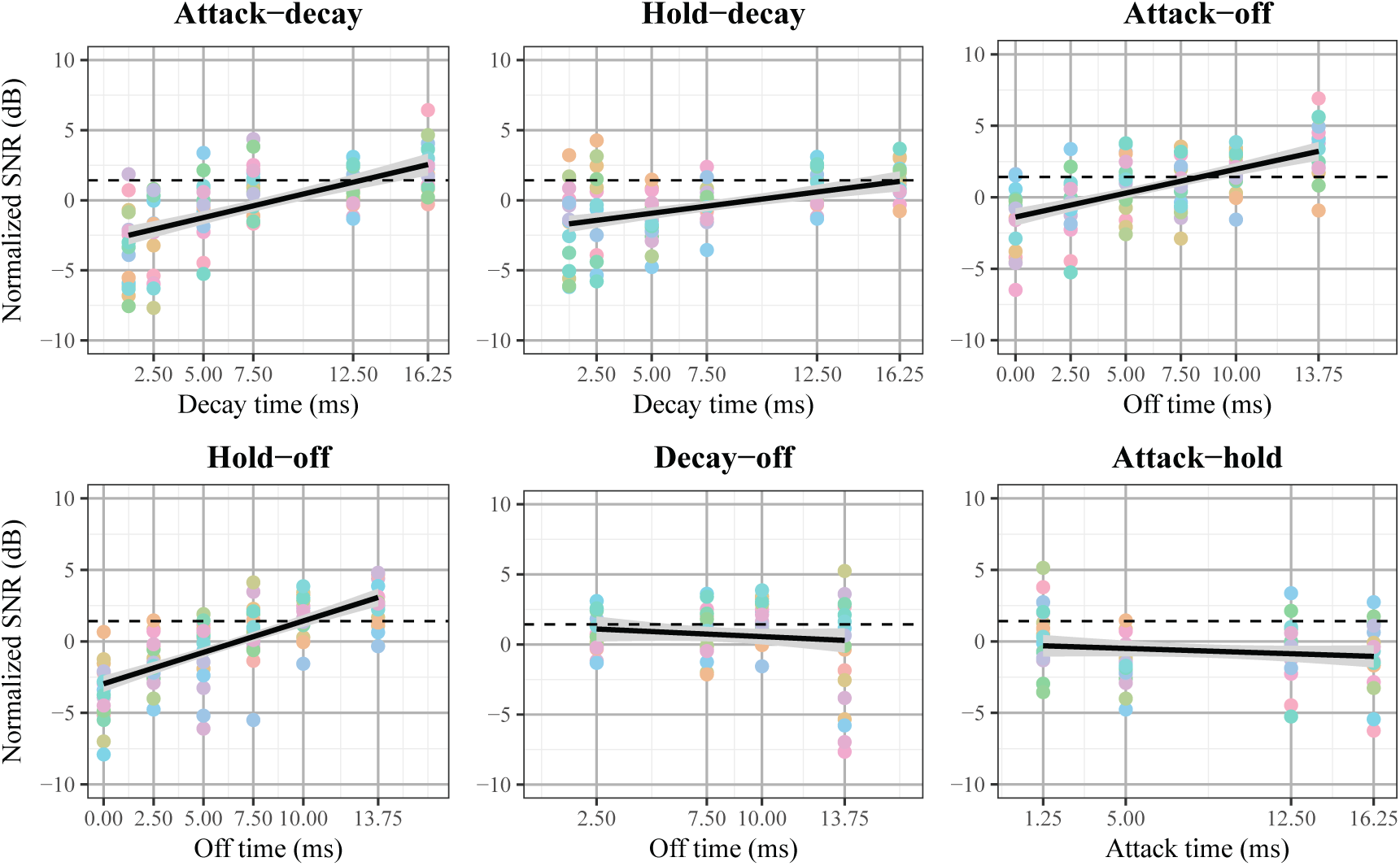
Normalized recording SNRs for the different stimuli and all subjects organized per condition. Data points of the same subject are represented in the same color. A linear regression line and its 95% confidence interval (grey) are plotted for each condition. The title of the subplot indicates the two envelope parameters that were covaried. The median normalized response SNR for the SAM stimulus is plotted as a dashed line (for further discussion on the comparison with SAM, see section 6.7).

For four conditions, linear regression showed a significant positive effect. Condition attack-decay showed an increase in response SNR for stimulus envelopes with larger decay time and smaller attack time (*β* = 0.35, t = 8.21, df = 112, p < 0.001). Similarly, condition hold-decay showed an increase in response SNR with longer decay and shorter hold time (*β* = 0.20, t = 5.66, df = 108, p < 0.001). For condition attack-off SNR increased with greater off time and smaller attack time (*β* = 0.33, t = 8.68, df = 115, p < 0.001), and finally condition hold-off showed larger SNRs for longer off time and shorter hold time (*β* = 0.44, t = 12.08, df = 124, p < 0.001). Note that it is hard to determine to what extent each of the two covaried parameters contributed to the observed effects.

In the remaining two conditions, i.e. decay-off (*β* = −0.12, t = −1.68, df = 81, p = 0.097) and attack-hold (*β* = −0.05, t = −1.21, df = 78, p = 0.23), the linear regression indicated no significant effect. This could indicate that neither of the two envelope parameters affected the response SNR. However, if this were the case for both of these conditions, it would mean that none of the four envelope parameters had an effect, which contradicts the results of the other conditions. Therefore, the flat curve in at least one of the two conditions had to result from two effects cancelling each other out. This left three possible scenarios to explain the observed results. The first was that shorter attack and hold time led to increased SNRs and off and decay time had no effect. The second option was the opposite: longer decay and off time resulted in larger response SNRs and attack and hold time had no effect. The final scenario was that all four factors had an effect: both shorter attack and hold time and longer decay and off time were related to larger SNRs.

Generalized linear mixed models with different fixed effects were constructed based on the scenarios specified above. The first model had attack and hold time as fixed effects (model attack-hold). Both effects were significant: attack time (t = −11.16, df = 454, p < 0.001) and hold time (t = −11.13, df = 454, p < 0.001). The second model had decay and off time as fixed effects (model decay-off) and these were also significant: decay time (t = 11.13, df = 454, p < 0.001) and off time (t = 10.12, df = 454, p < 0.001). The third scenario could not be translated to four fixed effects as this would lead to perfect multicollinearity (due to the covarying study design). However, we tested models with all possible combinations of three fixed effects and the third effect was never significant. Consequently, we concluded that the results are most likely explained by one of the first two models.

Model attack-hold and decay-off led to highly similar residual variance: 2.448 for decay-off and 2.452 for attack-hold, meaning they were equally good at explaining the data. This ambiguity can be expected from the covarying study design. Even when two parameters do not actually explain the results, the corresponding model will be significant because they covary with the actual predictors (the other two parameters). Despite that, it is still possible to distinguish the real model from the ‘false positive’ model. Since the two ‘false positive’ predictors each covary with both of the actual predictors, both ‘false positive’ effects will be a combination of the ‘actual’ effects. This leads to a large correlation between them. For that reason, the model with lower correlations between the fixed effects is more likely true. In our case, the correlation between effects is 0.433 for model attack-hold and 0.288 for model decay-off. We therefore conclude that changes in decay time and off time are the most likely cause of the observed changes in response SNR. More specifically, a 1 ms increase in decay time enhanced response SNR with 0.33 dB and a 1 ms increase in off time enhanced response SNR with 0.29 dB.

### 3.2. Response phase delay

The mean phase delay across all conditions and subjects was 342.8°. Mean phase delays per subject varied between 303° and 405° illustrating the inter-individual variance and the need for normalization. It is important to remark that we do not have enough information to interpret these phase delay values in an absolute way, but the aim of the study only requires relative comparison. Figure 4 shows normalized phase delay values with a regression line fitted for the different conditions. The three conditions where attack time was varied, showed an increase in phase delay with increasing attack time. Moreover, condition off-hold revealed a decrease in phase delay with increased off time. A generalize mixed model with random intercept shows that both attack time (t = 14.546, df = 454, p < 0.001) and off time (t = −2.278, df = 454, p = 0.023) were significant predictors of phase delay. A 1 ms increase in attack time was related to a raise in phase delay of 4.78 degrees. A 1 ms increase in off time led to a decrease in phase delay of 0.8 degrees.

**Figure 4:**
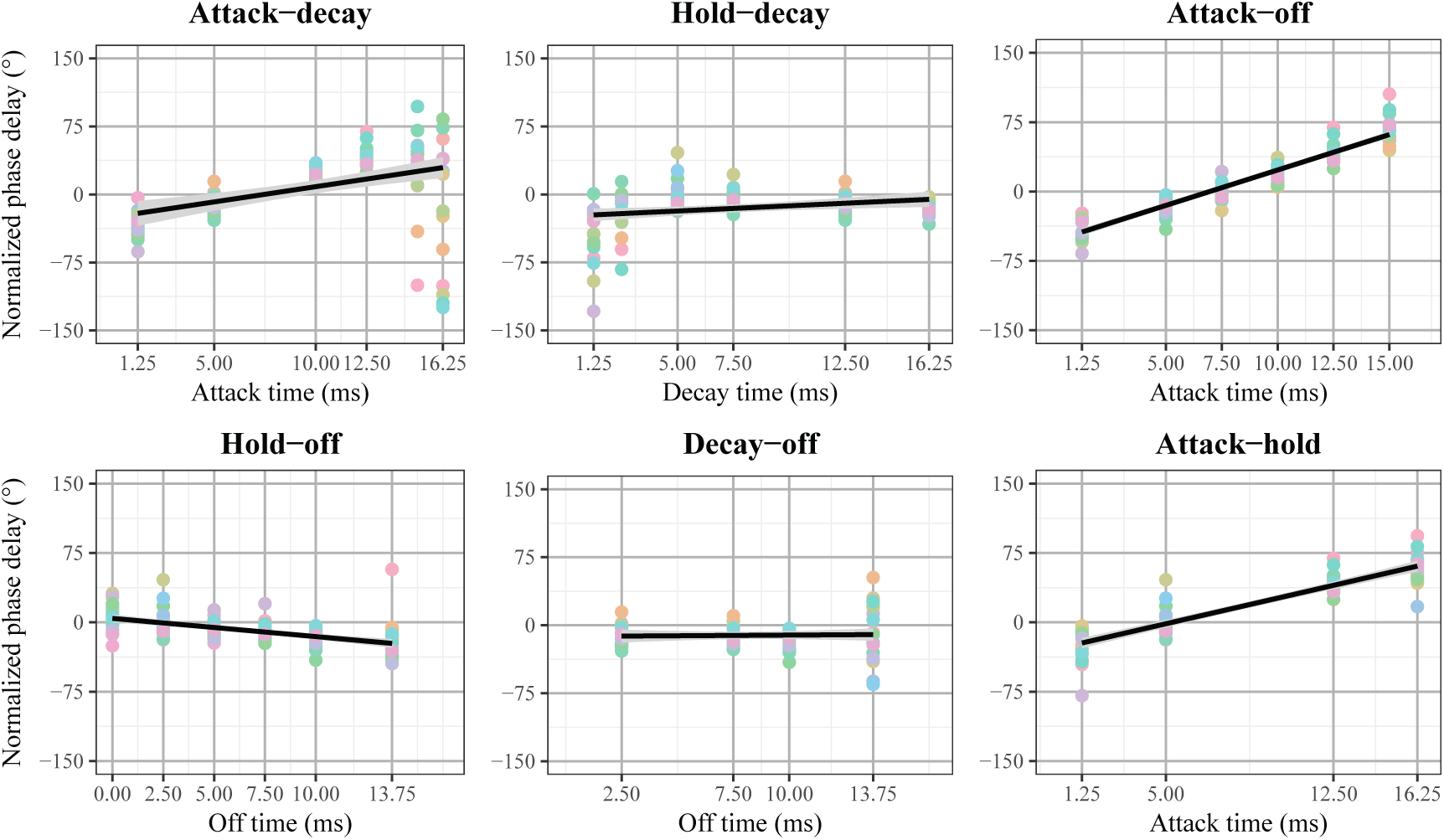
Normalized phase delay for the different stimuli and all subjects organized per condition. Data points of the same subject are represented in the same color. A regression line and its 95% confidence interval (grey) are plotted for every condition. The title of the subplot indicates the two envelope parameters that were covaried.

## 4. Model of the auditory periphery - methods

To help understand the effects of the envelope parameters found in the ASSR measurements, we simulated auditory nerve responses with a model of the auditory periphery by Bruce et al. (2018). This phenomenological model has a long history and was very recently updated (Carney, 1993; Zhang et al., 2001; Bruce et al., 2003; Zilany and Bruce, 2006, 2007; Zilany et al., 2009, 2014; Bruce et al., 2018). It requires a sound-pressure wave as input and provides simulated spike times from auditory nerve fibers with different characteristic frequencies (CF) as output. In this study, we used 60 different CFs logarithmically spaced between 250 and 8000 Hz. For every CF, the model simulated 50 auditory nerve fibers of which respectively 30, 10 and 10 fibers had low (0.1/s), median (4/s) and high (70/s) spontaneous firing rates. This means we simulated a total neural population of 3000 auditory nerve fibers with varying CFs and spontaneous firing rates. The output from the model can be visualized in a neurogram (an example is given in figure 5). For a more detailed description of the model, we refer to Bruce et al. (2018).

**Figure 5:**
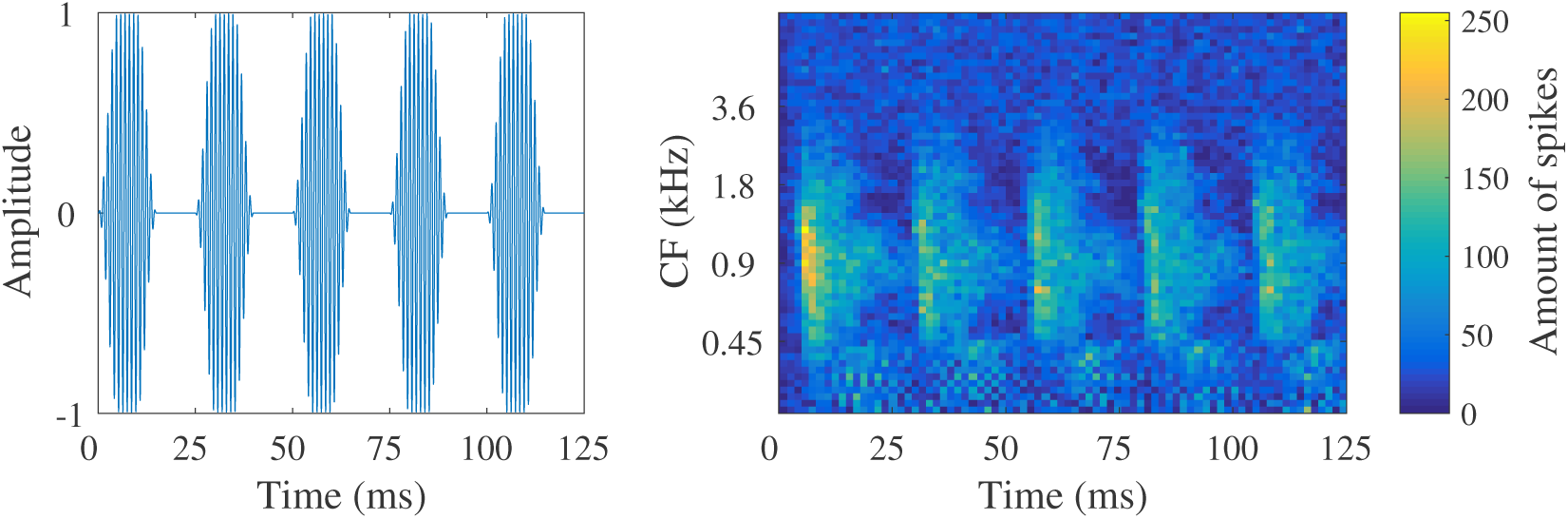
**Left:** 5 periods of the 20 period long stimulus used as input for the model. **Right:** corresponding output of the model of the auditory periphery by Bruce et al. (2018) visualized in a neurogram. Each time bin of the neurogram has a length of 3.2 ms. Note that the first period contains a response to the onset of the sound stimulus.

Auditory nerve responses were modelled for two different stimulus sets. The first stimulus sets was identical to the one for the ASSR measurements, but only 20 periods of each stimulus were used. The simulations for these stimuli provided insight in the relevancy of the simulations of the auditory periphery for the ASSRs, which are generated higher up in the auditory system. In the second stimulus set, each of the envelope parameters was altered independently, which means modulation frequency varied across stimuli. As explained before, this was avoided for the ASSR measurements because ASSR amplitude is heavily influenced by modulation frequency. However, since the model only regards neural responses in the auditory periphery, it is not dependent on modulation frequency. Consequently, the model allowed to study the independent effect of each of the envelope parameters in a way that was not possible with ASSRs. To construct this second stimulus set, one envelope parameter was varied between 1.25 and 51.25 ms and the others remained constant. When constant, decay and off time were equal to 5 ms and attack time was equal to 1.25 ms. Hold time had a constant value of 5 ms when decay or off time were varied and a constant value of 1 ms when attack time was varied. Modulation frequency ranged between 16 and 80 Hz and larger values of the envelope parameter were related to lower modulation frequencies.

ASSRs are generated by the total neural activity of a large population of neurons. Therefore, population responses were constructed from the model output by summing the spike counts together over all CFs (i.e. summing along the y-axis of the neurogram). This results in a waveform that reflects how the total firing rate at the level of the auditory nerve changes over time in response to the stimulus. The total waveform (20 periods) was averaged to obtain one simulated response period.. From the average response period, the minimum firing rate, maximum firing rate and neural modulation depth for the different envelope shapes was determined.

## 5. Model of the auditory periphery - results

### 5.1. Simulations for the ASSR stimuli

In figure 6, the simulated population response for each of the ASSR stimuli is shown. The simulated responses followed the periodicity of the input stimulus, with different response shapes for different envelope shapes. For instance, the width of the response peak was related to the broadness of the envelope shape, controlled by hold time. Furthermore, the time at which the neural response reached its maximum reflected the timing of the maximum amplitude in the envelope, controlled by attack time.

**Figure 6:**
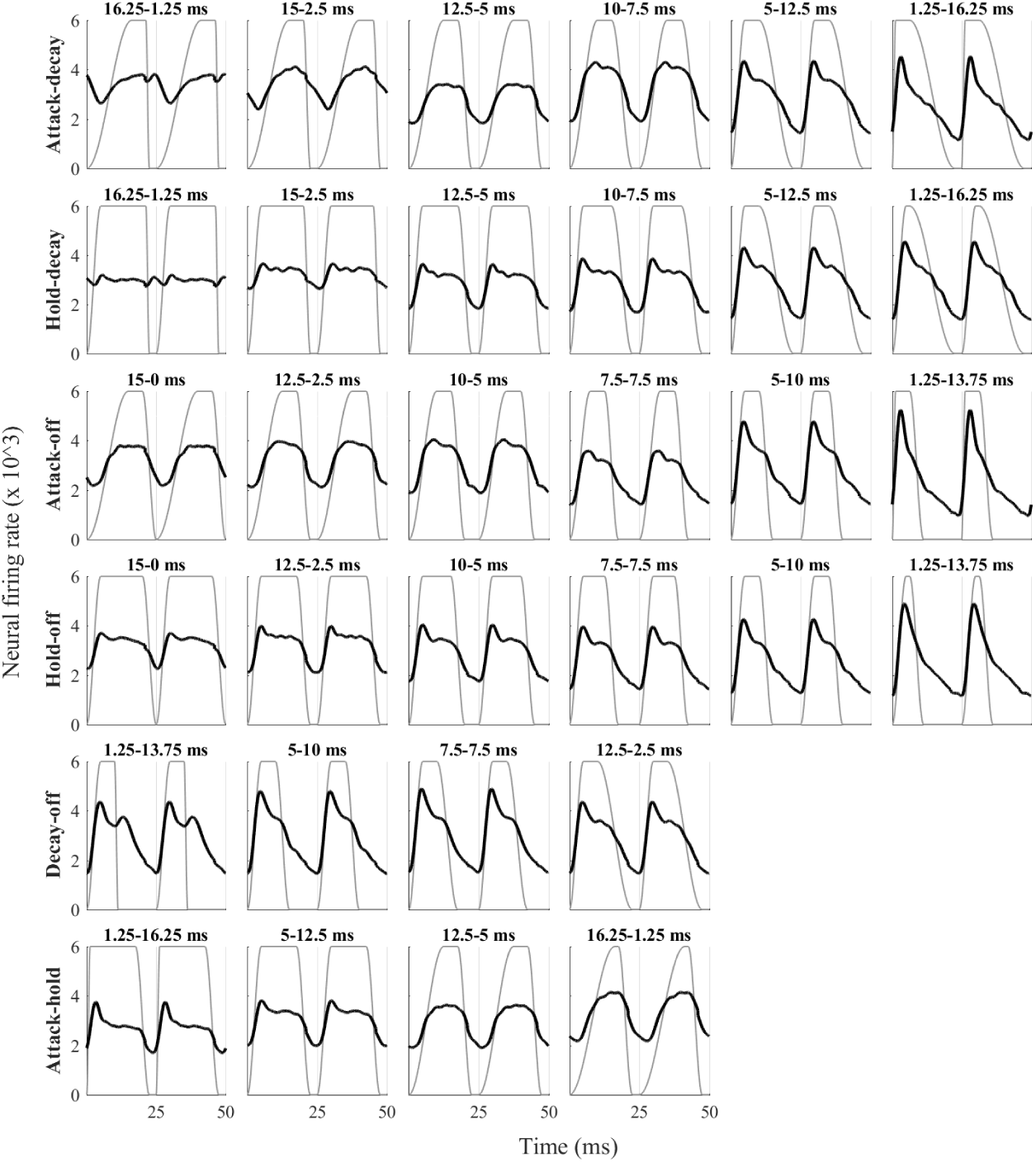
Simulated population responses for the ASSR stimuli based on the model of the auditory periphery by Bruce et al. (2018). The averaged period responses are repeated, such that two periods are visualized. For every simulated response, the envelope of the stimulus used as input for the model is plotted in grey. In the title of each subfigure, the values of the varied envelope parameters are written in the order they appear in the condition name.

There were also clear differences in the modulation depth of the responses. The modulation depths of the simulated responses, calculated as the difference between the maximum and minimum neural firing rate reached in a response period, are presented in figure 7. Linear regression revealed significant effects of envelope shape on response modulation depth in conditions attack-decay (*β* = 112, t = 10.09, df = 118, p < 0.001), hold-decay (*β* = 152.37, t = 10.142, df = 112, p < 0.001), attack-off (*β* = 172.89, t = 13.11, df = 118, p < 0.001) and hold-off (*β* = 160.9, t = 10.83, df = 118, p < 0.001). Conditions attack-hold (*β* = 11.17, t = 0.781, df = 78, p = 0.437) and decay-off (*β* = −1.984, t = −0.105, df = 78, p = 0.917) contained no significant effect. Just like for the ASSR results, it is hard to determine independent effects of the envelope parameters based on these results due to the covarying stimulus design.

**Figure 7:**
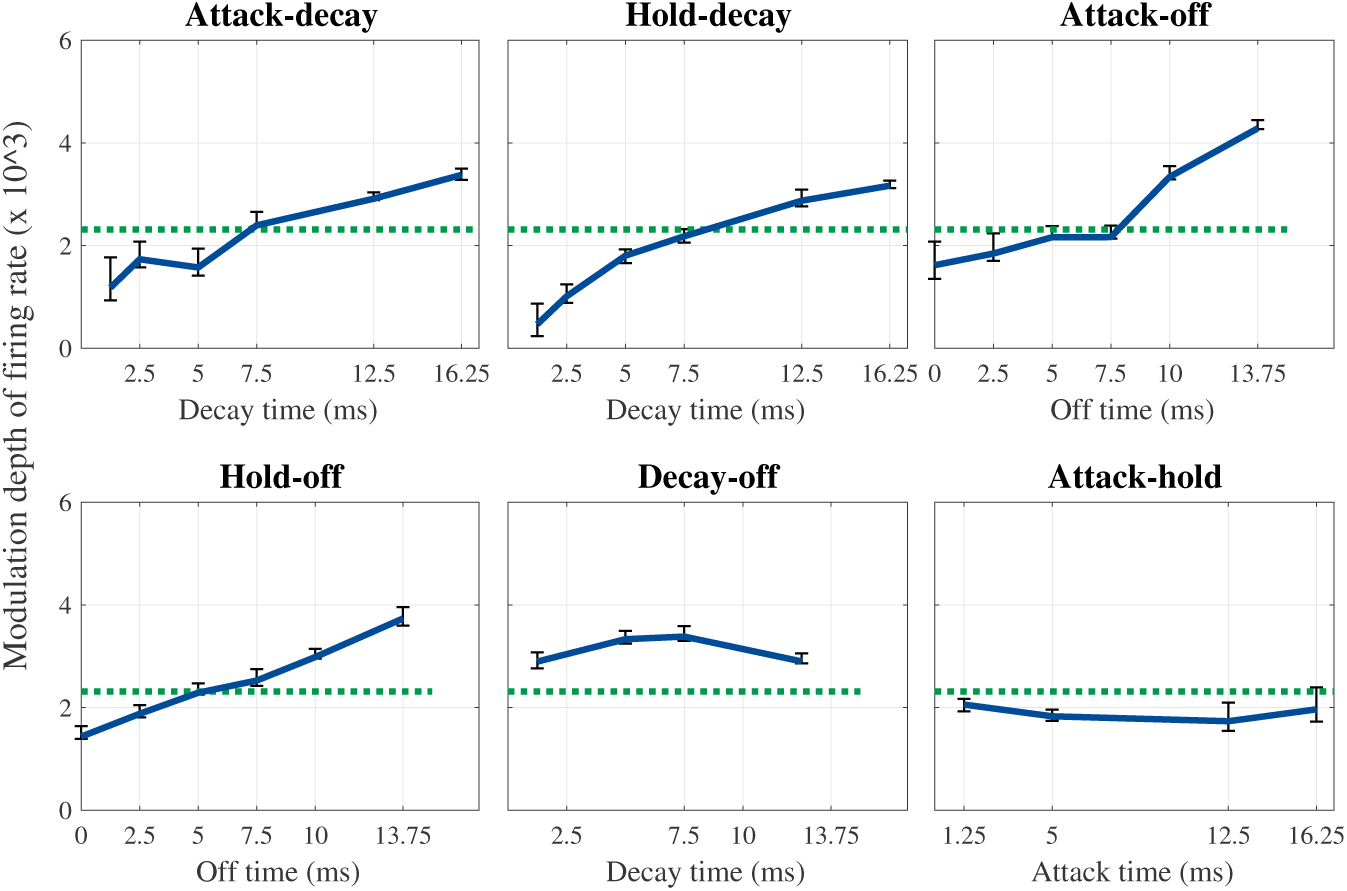
Modulation depth in function of envelope shape parameters for the different conditions. Variance over periods, which was calculated by the interquartile range, is plotted as errorbars. The simulated modulation depth for the SAM stimulus was 2312.9 (dotted line).

The results of the model were similar to the ones of the ASSR measurements (compare with figure 3). To quantify the resemblance, we calculated the Pearson correlation between the simulated modulation depths and the measured response SNRs. Since the model was designed to predict the average case, we correlated the simulated modulation depth with the median response SNR across subjects. The relation between the two variables is shown in figure 8. The correlation coefficient was equal to 0.812 (t = 6.678, df = 23, p < 0.001), showing that the response SNR was strongly related to the modulation depth of the population response at the level of the auditory nerve. It also confirmed that simulations of the response at the auditory periphery using the model of Bruce et al. (2018) are a reliable way to study and interpret changes in ASSRs related to envelope shape.

**Figure 8:**
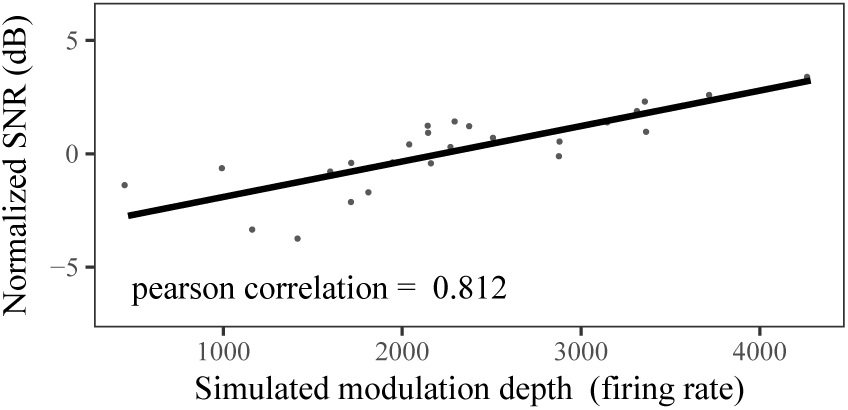
Correlation of the modulation depth of the neural response with the median normalized recordings SNR across subjects.

### 5.2. Simulations for the stimuli with varying modulation frequency

Figure 9 shows the population responses for a subset of the second stimulus set where each of the envelope parameters was varied independently. Figure 10 presents the simulated modulation depths for the complete stimulus set. Both figures show that changing the duration of the attack or hold, had almost no influence on the modulation depth of the response. On the other hand, modulation depths increased logarithmically when decay and off time were increased from 1.25 ms to 28.75 ms. Extending decay or off time beyond this range added little benefit.

**Figure 9:**
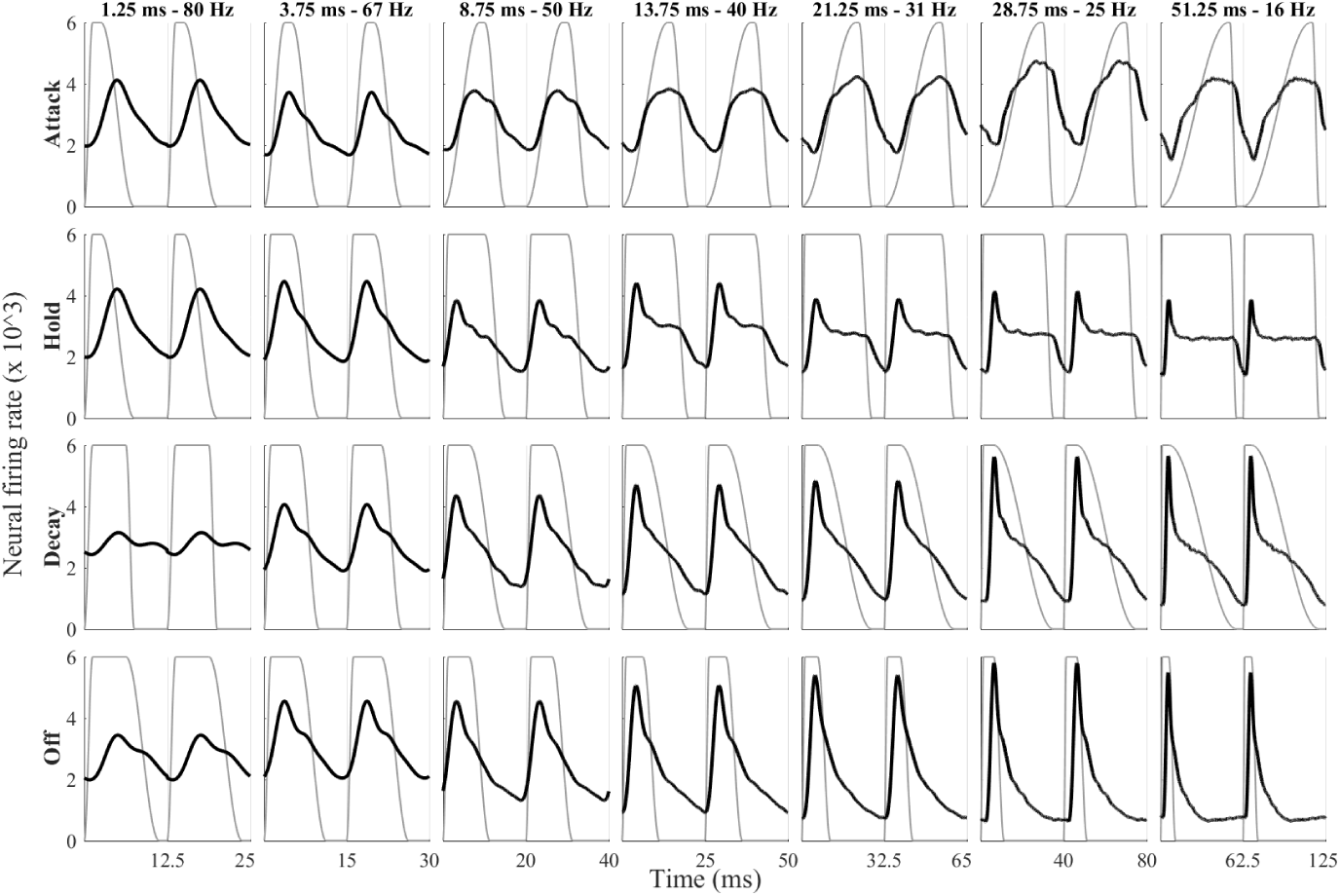
Simulated population responses for a subset of the second stimulus set based on the neurograms outputted by the model of the auditory periphery by Bruce et al. (2018). The full stimulus set contained 11 stimuli per condition, corresponding to 11 different values of the varied parameter. In this figure, population responses are shown for only 7 stimuli per condition, but they were chosen such that the full range was represented. The row labels indicate which envelope parameter was varied. The column labels show the value of the varied envelope parameter and the modulation frequency for the stimuli in that column. Note that the scale of the x axis differs along each row.

**Figure 10:**
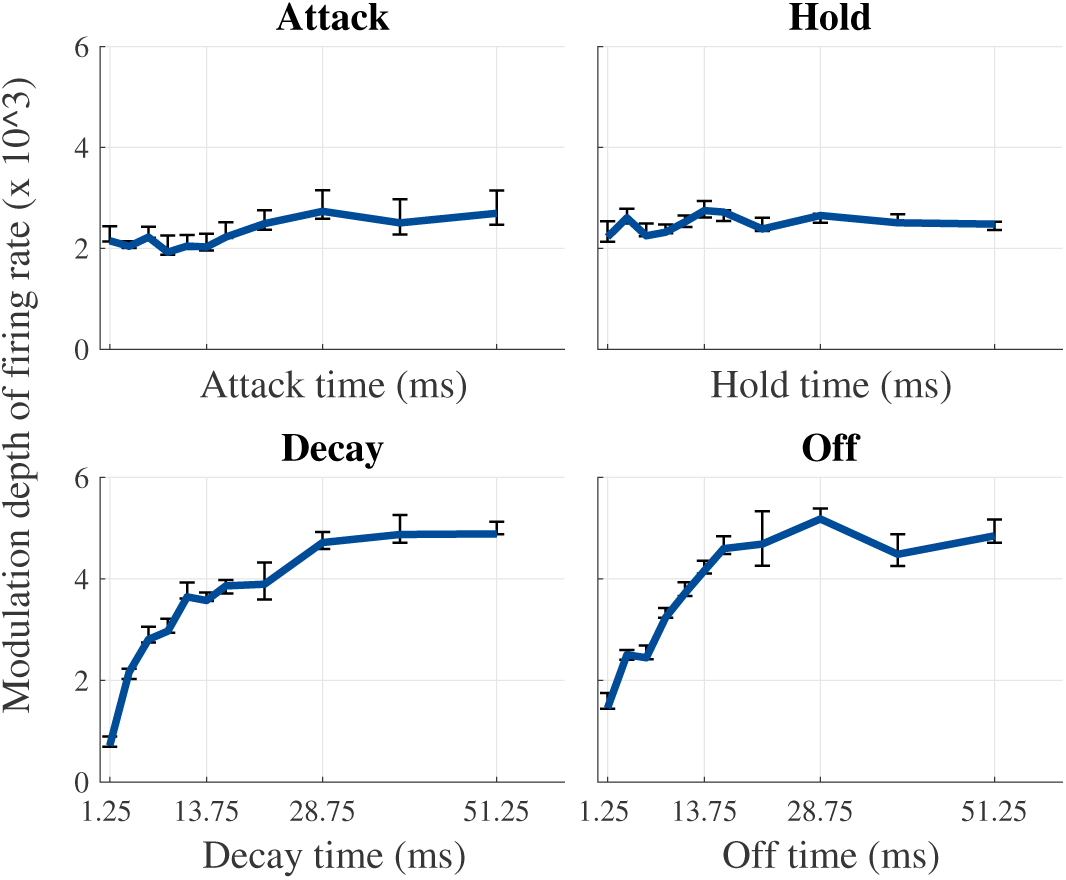
Modulation depth in function of envelope shape parameters for the different conditions. Variance over periods, which was calculated by the interquartile range, is plotted as errorbars.

Since modulation depth of the neural population response is highly correlated with ASSR SNR, it makes sense to further investigate the two factors contributing to modulation depth, i.e. minimum and maximum firing rate. Figure 11 shows the minimum and maximum firing rates of the simulated population responses for the different conditions. Minimum and maximum rate were minimally affected by attack and hold time. Minimum firing rate decreased logarithmically for increasing decay and off time. Similarly, maximum firing rate increased logarithmically for increasing decay and off time. We conclude that both changes in minimum firing rate and maximum firing rate contributed to the effect of decay and off time on the modulation depth of the auditory nerve response.

**Figure 11:**
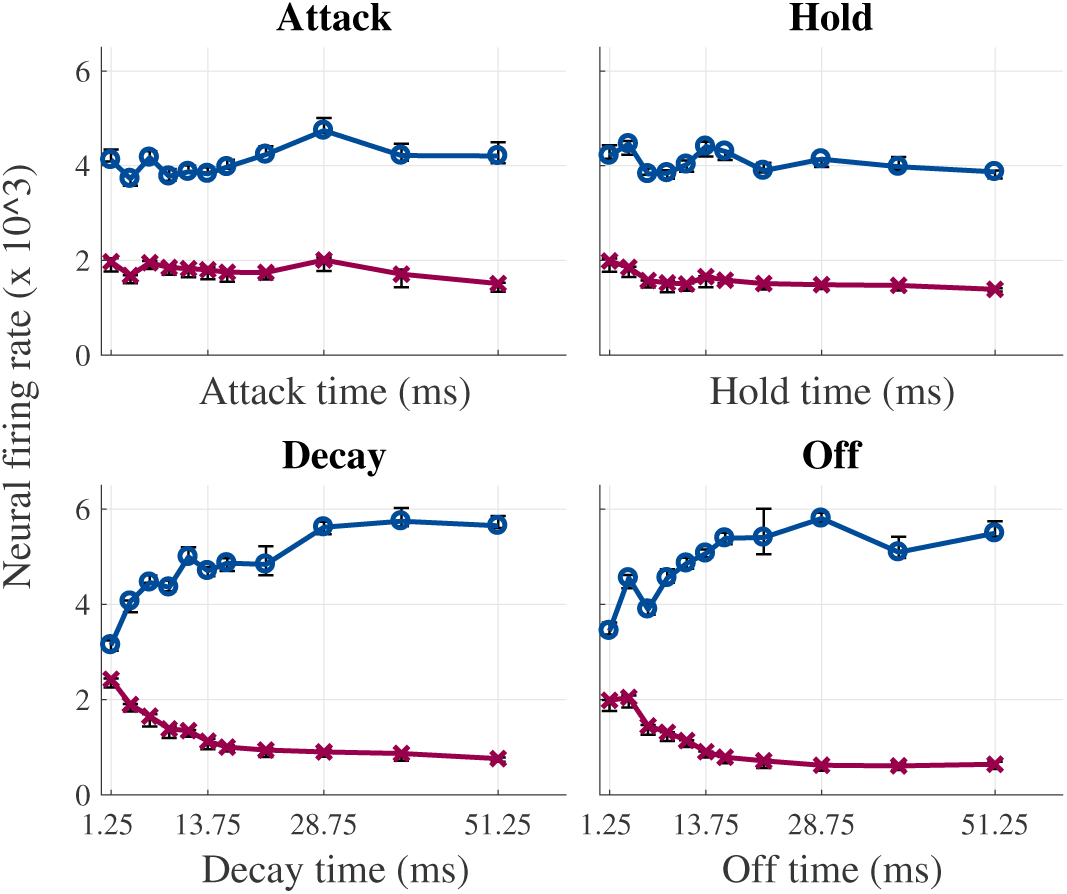
Minimum (x) and maximum firing rate (o) of the simulated population responses for the different conditions. Errorbars are plotted, but were generally around the same size as the markers used to represent the datapoints.

## 6. Discussion

The objective of this study was to investigate the effect of envelope shape on ASSRs. We studied how response SNR and phase delay are affected by the four envelope parameters: attack time, hold time, decay time and off time. Actual ASSR measurements were complemented with simulations of the auditory nerve response using the model of Bruce et al. (2018).

### 6.1. The effect of envelope shape on response SNR

ASSR measurements showed that response SNR was significantly influenced by envelope shape. The response pattern could be explained by three scenarios: an effect of attack time and hold time, an effect of decay time and off time or an effect of all four parameters. Based on the lower correlation of effects when modelled with a linear mixed model, we concluded that decay time and off time were most likely the dominant factors. The linear mixed model showed that a 3 ms increase in decay or off time led to a 1 dB increase in response SNR. The ASSR measurements alone did not allow to rule out that attack time and hold time had a (smaller) effect as well. Simulations of the auditory nerve response with the model of Bruce et al. (2018) allowed to vary each envelope parameter independently, as changes in modulation frequency had no confounding effect. The model results revealed a logarithmic effect of off and decay time on the modulation depth of the population response. Longer off and decay time were associated with an increase in modulation depth, because of both a higher maximum firing rate and a lower minimum firing rate. There was no effect of attack or hold time. Moreover, a large correlation between the modulation depth of responses modelled for the ASSR stimuli and the SNR of the measured ASSRs was found. Together, this confirms that ASSRs are affected by envelope shape, where longer decay and off time provide larger responses.

### 6.2. Physiological processes behind the effect of envelope shape on response SNR

To understand the physiological processes behind the effect of decay and off time on the SNR of the ASSR, it is valuable to look at its decomposition in the effect on minimum firing rate and the effect on maximum firing rate (see figure 11). The decrease in minimum firing rate with increasing off time is likely a direct consequence of the absence of stimulation. Longer off time allows neural activity to subside more, leading to a larger drop in firing rate. For off time longer than 28.75 ms, no further decrease in minimum firing rate was observed. This likely reflects that the minimum firing rate is approaching the spontaneous firing rate.

Decay time is important for minimum firing rate because it determines the steepness of the decay. Our results indicate that a slower, less steep decay provides a lower minimum firing rate. A long decay time will allow neural firing to subside already during the decay, such that during off time a lower minimum firing rate can be reached. In contrast, a short decay time creates a very sudden change in envelope slope, which elicits a large and spectrally broad burst in neural activity. The shorter the decay time, the stronger and broader the activity. This can be seen in the neurograms in figure 12 and also in figure 6 where a second response peak appears after short decays (e.g. in conditions attack-decay, hold-decay and decay-off). The burst in neural firing induced by short decay time counteracts the subsiding of neural firing rate. This way, a short decay can undo the positive effect of off time, as evidenced by figure 12. We can now also explain the relatively flat curve for modulation depth in condition decay-off in figure 7 and for SNR in figure 3. A long decay time will allow neural firing to subside already during the decay, allowing the neural response to die out to a lower minimum even if off time is very short. On the other hand, when off time is very long, minimum firing can still be reached with a very short decay, because the burst in neural activity will have died out before the end of the off time (see also condition decay-off in figure 6).

**Figure 12:**
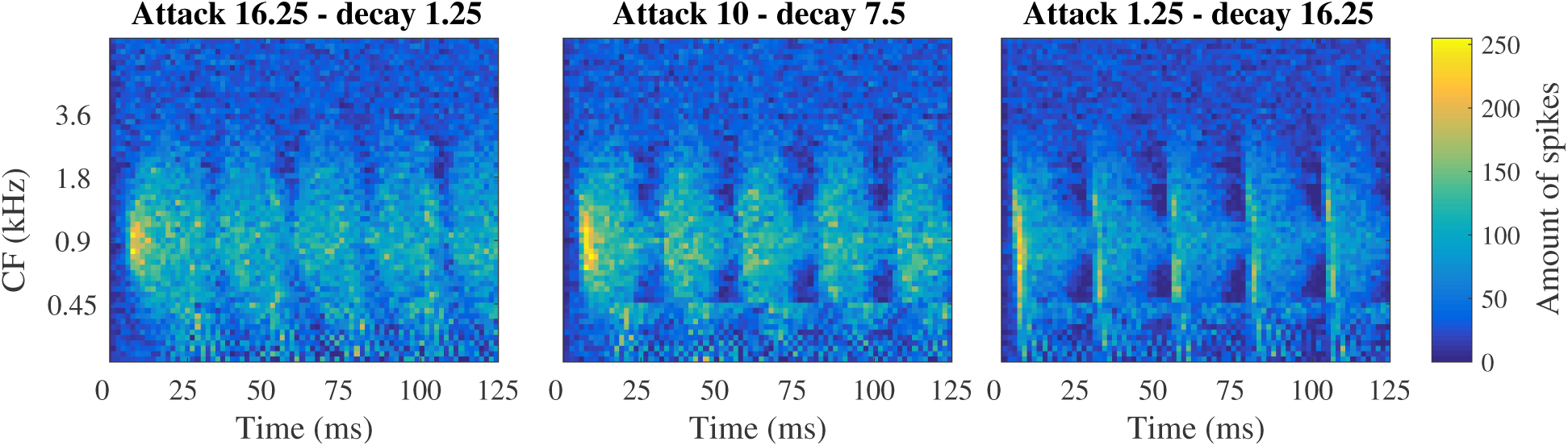
Neurograms of three responses in condition attack-decay

Based on the fact that neural firing probability is largest during the rising phase of the envelope period, one would expect maximum firing rate to be influenced mainly by attack time. This is reflected in the theory that shorter attack times (or steeper attack slopes) cause larger neural responses (e.g. John et al. (2002)). Indeed, when looking at figure 6, it can be seen that short attack times, i.e. 2.5 - 5 ms long, led to a sharp peak in the neural response. However, as can be observed in condition attack-hold, a higher maximum firing rate was reached with long attack time than with short attack time. Hence, a short attack time increased the peakedness of the population response but not the overall maximum firing rate. Similarly, in figure 11 it can be seen that short attack times did not provide larger maximum firing rates.

Figure 11 shows that maximum firing rate is influenced by decay and off time, i.e. the same factors that influenced minimum firing rate. It is likely that their effects on maximum firing rate are a consequence of their effect on minimum firing rate. A lower minimum firing rate means that less neurons are firing in the time period before each attack. This causes more neurons to be recuperated from their refractory period at the time of the next attack. Consequently, a larger amount of neurons is ready to fire in response to the attack and a larger maximum firing rate can be reached. In this way, decay and off time affect the modulation depth of the population response, and therefore response SNR, by both decreasing minimum firing rate and increasing maximum firing rate.

### 6.3. Comparison with other studies

John et al. (2002) found that exponential sine envelopes provide larger ASSR responses than SAM stimuli and suggested that this was caused by longer off time and shorter attack time. The current study confirms the positive effect of longer off time on the ASSR response, but found an effect of decay time instead of attack time. However, John et al. (2002) could not distinguish between the effects of attack and decay time, since taking an exponent of a sinusoidal envelope (N>1) changes the attack and decay in equal ways. John et al. (2002) assumed that changes in attack time explained the results, rather than decay time, as they believed the ASSR to be “an onset response - evoked primarily by the repeating rise-functions of the stimulus”. The results of the present study show that the falling phase of the stimulus envelope is at least equally (if not more) important.

The results found in this study also differ from the results of several studies concerning the effect of envelope shape on ITD perception (Bernstein and Trahiotis, 2002, 2009; Klein-Hennig et al., 2011; Laback et al., 2011). This discrepancy can most likely be explained by differences in the neural processes that underlie ASSRs and ITD perception. ASSRs are periodic responses to a periodic signal, for which the modulation depth of the neural response has proven to be essential. On the other hand, ITD perception is the detection of time differences between the sound input at the two ears. This detection process likely relies most heavily on the onset and attacks of the sound. It is therefore not surprising that the ITD perception studies found a significant effect of attack time (which was not found for ASSRs). Moreover, they also found a significant effect of off time. As we have shown, a longer off time decreases minimum firing rate, which increases firing capacity for the next attack - which likely facilitates the detection of time differences. Since decay time also influences maximum firing rate at the next attack, one would expect there to be a significant effect of decay time on ITD perception as well. The only study where decay time is investigated, i.e. Klein-Hennig et al. (2011), found that ITD perception was close to impossible based on time differences in the decay of the sound alone. They therefore concluded that decay time had no or only a supporting effect. Indeed, the results of the present study point towards a supporting effect of decay time on the perception of ITDs present at the attack - which was not investigated by Klein-Hennig et al. (2011).

Greenberg et al. (2017) performed single neuron recordings in the midbrain of guinea pigs to evaluate the importance of envelope shape for ITD perception at a neural level. Apart from neural ITD sensitivity, they also analysed neural phase-locking and temporal dispersion of neural firing in function of envelope shape. The results indicated stronger phase-locking and lower temporal dispersion for envelopes with longer off time, which is in agreement with the results of the present study. Besides this, they also reported stronger phase-locking and lower temporal dispersion for stimuli with shorter attack time, which is not in line with the present findings. However, the stimuli varied both in attack and decay time and the study neglected the possibility of an effect of decay time. In fact, both the stronger phase-locking and lower temporal dispersion can be explained equally well, if not better, by long decay time than by a short attack time.

### 6.4. The effect of envelope shape on phase delay

We also looked into the effect of envelope shape on the phase delay of the responses. Phase delay was significantly influenced by attack time, i.e. longer attack times led to larger phase delays. John et al. (2002) wrote that response latency (which is derived from phase delay) is related to changes in the timing of the maximum slope of the stimulus envelope. Since peak level was kept constant in this study, attack slope was directly related to attack time. Therefore the conclusion of the present study and the study of John et al. (2002) are in agreement. The ASSR measurements of the current study also revealed an effect of off time on phase delay: a longer off time led to a slight decrease in phase delay. This could reflect how longer off time allows neurons to recuperate more from their refractory periods supporting a faster response to the next attack.

### 6.5. The effect of presentation level

It could be argued that the high presentation level, i.e. 75 dB SPL, used in this study is the reason why we did not find an effect of attack time on the ASSR amplitude. It could be that the large presentation level saturated the neurons so that the highest neural firing rate was always reached and variations in maximum could not be fully observed. To investigate this possibility, we simulated responses for the same stimulus set but now presented at 40 dB SPL. This level was chosen in analogy with John et al. (2002), who presented their stimuli at a lower level since they were trying to optimize stimuli for objective measurement of hearing thresholds. Results showed that maximum firing rate was generally lower for the lower presentation level, but they did not reveal an effect of attack time.

As explained in the methods section, all stimuli were presented at the same peak level. This ensured that the slope of the attack depended only on attack time. Moreover, this avoid confounding effects as changes in peak levels may result in differences in the amount of neurons recruited (Liberman, 1978; Laback et al., 2011). A disadvantage of this approach is that it introduced differences in the presentation level across the stimuli. To evaluate whether this confounded the results, we registered the presentation level of each stimulus in dB SPL. A linear regression model showed that level was influenced by two of the four envelope factors, i.e. hold time (t = 10.36, df = 22, p < 0.001) and off time (t = −8.65, df = 22, p < 0.001). Longer hold times and shorter off times were related to higher presentation levels when peak level was kept constant. More in detail, an increase of about 1 dB SPL was observed when hold time was increased with 5 ms or off time was decreased with 5 ms. ASSR SNR is known to increase with higher stimulus presentation levels (Picton et al., 2003; Van Eeckhoutte et al., 2016), so if level difference would explain the observed results, there should be larger ASSRs for longer hold times and shorter off times. However, there was no significant effect of hold time and the effect of off time occurred in the opposite direction. Therefore we conclude that presentation level did not cause the observed effect of envelope shape on the ASSR.

### 6.6. Relevance of the simulated auditory nerve responses to ASSRs

The simulations in this study were done with a model of the auditory periphery. The fact that the modulation depth of the simulated responses correlated highly with the measured response SNRs, shows that the simulations are highly relevant for ASSRs. This makes the model of Bruce et al. (2018) of great value for research on ASSRs. For example, the model provides a tool to explore a wide range of stimulus parameters in a fast way before choosing parameters to use in actual measurements. Moreover, the success of the simulations also implies that ASSR characteristics are determined for a surprisingly large part by processing in the auditory periphery. Naturally, an even larger similarity between simulations and measured ASSRs is expected when subcortical and cortical processing are also included in the model. An example of such a model is the model by Verhulst et al. (2018) which includes processing up to the inferior colliculus.

### 6.7. Implications of this research

ASSRs are clinically used to objectively measure hearing thresholds. In clinical practice, response measurement should be optimized as much as possible. Stimuli that provide larger responses can decrease measurement time and sensitivity to subject-related and environmental noise. The results of this research indicate that the largest response SNRs are obtained when there is sufficient time with no stimulation during the falling phase of the envelope period. Moreover, sharp changes in envelope slope during the falling phase, like a sharp decay, should be avoided.

The most commonly used stimulus to evoke ASSRs is the SAM stimulus. It has a long decay time (1/2 of the period), but no off time. When comparing the response SNR (see figure 3) or simulated modulation depth (see figure 7) for a SAM stimulus to those for the other stimuli, it can be seen that stimuli with relatively long decay and/or off time indeed provided better results than the SAM stimuli. The results of this research point towards the most optimal stimulus having a short attack time (1.25 ms), short hold time and the rest of the period duration roughly divided between decay and off time. One of the stimuli used in condition attack-off comes close to this optimum: attack = 1.25 ms, hold = 5 ms, decay = 5 ms and off = 13.75 ms. The median gain in ASSR SNR over subjects for this stimulus compared to the SAM stimulus was 1.63 dB (interquartile range = 0.11-3.17 dB). In terms of response amplitude there was a median gain of 89 nV (interquartile range = 0-140 nV) which corresponds to an enhancement of 27% (interquartile range = 0-40 %). However, this conclusion is specific to 40 Hz modulation. For higher modulation frequencies (e.g. 80 Hz, which is often used in clinical practice), only very short decay and off time are possible due to short period duration, decreasing the potential benefit. The opposite is true for lower modulation frequencies, where larger benefits are possible.

## 7. Acknowledgements and Funding

Authors would like to thank all subjects who participated in the study. We also want to acknowledge Stephanie Nelis for her help with data collection. This research was funded by TBM-project LUISTER (T002216N) from the Research Foundation Flanders (FWO) and also jointly by Cochlear Ltd. and Flanders Innovation & Entrepreneurship (formerly IWT), project 50432. Additionally, this project has received funding from the European Research Council (ERC) under the European Unions Horizon 2020 research and innovation programme (grant agreement No. 637424, ERC starting grant to Tom Francart). The first author, Jana Van Canneyt, is supported by a PhD grant for Strategic Basic research by the Research Foundation Flanders (FWO), project number 1S83618N.

## 9. Abbreviations

ASSR: auditory steady-state response
CF: charactertistic frequency
EEG: electroencephalogram
ITD: interaural time difference
SAM: sinusoidal amplitude modulation
SNR: signal-to-noise ratio

